# Remdesivir, Molnupiravir and Nirmatrelvir remain active against SARS-CoV-2 Omicron and other variants of concern

**DOI:** 10.1101/2021.12.27.474275

**Authors:** Laura Vangeel, Winston Chiu, Steven De Jonghe, Piet Maes, Bram Slechten, Joren Raymenants, Emmanuel André, Pieter Leyssen, Johan Neyts, Dirk Jochmans

**Affiliations:** KU Leuven, Department of Microbiology, Immunology and Transplantation, Rega Institute, Laboratory of Virology and Chemotherapy, Leuven, Belgium; KU Leuven, Department of Microbiology, Immunology and Transplantation, Rega Institute, Laboratory of Clinical and Epidemiological Virology, Leuven, Belgium; University Hospitals Leuven, Department of Laboratory Medicine, Leuven, Belgium; KU Leuven, Department of Microbiology, Immunology and Transplantation, Rega Institute, Laboratory of Clinical Bacteriology and Mycology, Leuven, Belgium

## Abstract

We assessed the *in vitro* antiviral activity of remdesivir and its parent nucleoside GS-441524, molnupiravir and its parent nucleoside EIDD-1931 and the viral protease inhibitor nirmatrelvir against the ancestral SARS-CoV2 strain and the five variants of concern including Omicron. VeroE6-GFP cells were pre-treated overnight with serial dilutions of the compounds before infection. The GFP signal was determined by high-content imaging on day 4 post-infection. All molecules have equipotent antiviral activity against the ancestral virus and the VOCs Alpha, Beta, Gamma, Delta and Omicron. These findings are in line with the observation that the target proteins of these antivirals (respectively the viral RNA dependent RNA polymerase and the viral main protease Mpro) are highly conserved.

## Main Text

One and a half year after the start of the global COVID-19 pandemic caused by severe acute respiratory syndrome coronavirus 2 (SARS-CoV-2), multiple variants have emerged. These can be harmless or slightly beneficial for the virus, causing for example increased transmission, virulence, or immune escape [1-3]. SARS-CoV-2 genetic diversification was initially considered slow when the virus was spreading in early 2020. The first official *variant*, a single spike D614G mutation found in early European lineages, was linked to more efficient transmission [4] and rapidly spread to become the dominant viral strain worldwide. Late 2020, multiple variants emerged that spiked regional epidemics. Five ‘variants of concern’ (VOC) have been identified (Alpha, Beta, Gamma, Delta and Omicron). All have characteristic mutations (www.ecdc.europa.eu/en/covid-19/variants-concern). The spike (S) glycoprotein appears especially prone to accumulate mutations [5] and all of the circulating VOCs have some mutations that favor evasion from the host immune response [6]. Numerous spike-protein based vaccines were developed and vaccination programs are running at full speed. However, studies of sera and emerging real-world evidence indicate that Omicron escapes the immunity whether from previous infection or vaccination [7].

Several direct-acting antivirals against SARS-CoV-2 have been approved or are advancing in clinical development. They can be divided in two classes, monoclonal antibodies (mAbs) directed against the Spike protein and small molecules interfering with the viral replication machinery. mAbs are administered intravenously but studies are underway to explore intramuscular or subcutaneous administration which would overcome the requirement of a hospital setting for dosing. [8]. Recent cell culture data indicates that the SARS-CoV-2 variant of concern (VOC) Omicron is not susceptible to most of the approved mAbs making it unlikely that their clinical efficacy will be maintained [9, 10].

The direct-acting small-molecule SARS-CoV-2 antivirals that have received approval or emergency use authorization do not target the variable spike-protein but target either the conserved viral RNA-dependent RNA polymerase (RdRp) or the conserved viral main protease (Mpro or 3CL protease). Remdesivir, a monophosphoramidate prodrug of the nucleoside GS-441524, originally developed to treat Ebola virus infections, inhibits the RdRp of SARS-CoV-2. It was the first antiviral approved or authorized for emergency use to treat COVID⍰19 in several countries. Remdesivir improves clinical outcomes in patients hospitalized with moderate-to-severe disease and it prevents disease progression in outpatients [11, 12]. While remdesivir requires intravenous administration, an oral prodrug of GS-441524 is being developed [13]. Molnupiravir (MK-4482 or EIDD-2801), a prodrug of the nucleoside analogue EIDD-1931 (β-D-N4-hydroxycytidine), is another inhibitor of the viral RdRp and was originally developed against different RNA viruses such as influenza [14]. A phase 2a clinical trial of molnupiravir in patients with COVID-19 shows accelerated SARS-CoV-2 RNA clearance and elimination of infectious virus [15]. This orally bioavailable drug was recently authorized in the UK for use in people who have mild to moderate COVID-19 and who have at least one risk factor for developing severe illness. Also, the U.S. FDA issued an emergency use authorization (EUA) in infected adults who are at high risk for progression to severe COVID-19, and for whom alternative COVID-19 treatment options are not accessible or clinically appropriate.

Another target for antiviral drugs is the viral main protease Mpro (or 3CL protease), a cysteine protease which cleaves the two polyproteins (pp1a and pp1ab) of SARS-CoV-2 at multiple locations, resulting in the various non-structural proteins, which are key for viral replication. Nirmatrelvir (PF-07321332), is an irreversible inhibitor of SARS-CoV-2 Mpro that is co-formulated with ritonavir allowing an oral route of administration (known as Paxlovid). When treatment is initiated during the first days after symptom onset, it results in roughly a 90% protection against severe COVID-19 and hospitalization [16]. Even though the Mpro-gene can be slightly affected by evolutionary mutations, the antiviral potency does not seem to be compromised [17].

We here assess the *in vitro* antiviral effect of GS-441524, remdesivir, EIDD-1931, molnupiravir and nirmatrelvir against the various SARS-CoV-2 VOCs, including Omicron.

The SARS-CoV-2 antiviral assay is based on the previously established SARS-CoV assay[18]. Upon infection the fluorescence of VeroE6-eGFP cell cultures declines due to a cytopathogenic effect. In the presence of an antiviral compound, the cytopathogenicity is inhibited and the fluorescent signal maintained. To this end VeroE6-GFP cells (kindly provided by Marnix Van Loock, Janssen Pharmaceutica, Beerse, Belgium), were used as described previously [19, 20]. Since VeroE6 cells show a high efflux of some chemotypes [21], the antiviral assays were performed in the presence of the P-glycoprotein (Pgp) efflux inhibitor CP-100356 (0.5 μM). A SARS-CoV-2 strain grown from the first Belgian patient sample (GHB-03021/2020), was used as ancestral strain as it is closely related to the prototypic Wuhan-Hu-1 2019-nCoV (GenBank accession number MN908947.3) [22]. All the other isolates were obtained from patients in Belgium and more information can be found in GISAID (Alpha = EPI_ISL_791333; Beta = EPI_ISL_896474; Gamma = EPI_ISL_1091366; Delta = EPI_ISL_2425097; Omicron = EPI_ISL_6794907). The multiplicity of infection (MOI) was kept constant for the different VOC to allow comparison of the potency.

Our *in vitro* results show that GS-441524, remdesivir, EIDD-1931, molnupiravir and nirmatrelvir retain their activity against all current VOCs including Omicron (Figure 1). The maximal change of the median EC_50_s over the different VOC is for each compound < 3x (1.8x for GS-441524, 1.6x for remdesivir, 2.5x for the series EIDD-1931 and molnupiravir and 2.5x for nirmatrelvir). In our experience these results are within the normal range of measurement error. For example, the ratio between the 95% and 5% percentile of the EC_50_s of GS-441524 on GHB-03021/2020 is 2.9x (n=206; calculated using Graphpad v9.2.0). So only differences in EC_50_s of >3x and with statistical significance should be considered as meaningful differences using this methodology. The individual EC50s values of this study are available at Mendeley Data (doi: 10.17632/bmjc74dyjs.1).

**Figure 1:**
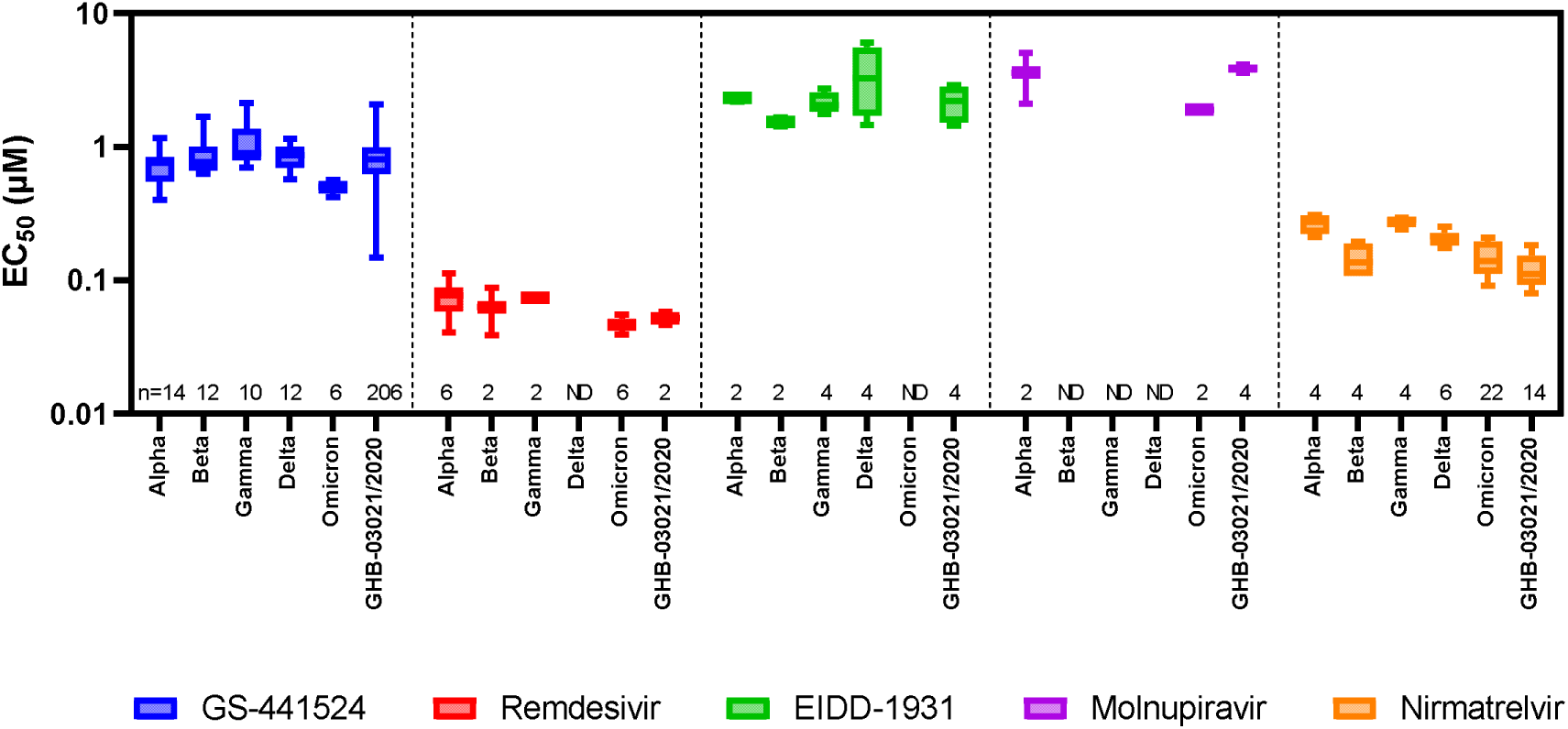
Activity of various antivirals upon infection of VeroE6-GFP cells with different SARS-CoV-2 VOC. VeroE6-GFP cells were pre-treated overnight with serial dilutions of the compounds. The next day, cells were infected with SARS-CoV-2 at a multiplicity of infection (MOI) of 0.001 tissue culture infectious dose (TCID_50_) per cell. The number of fluorescent pixels of GFP signal, determined by high-content imaging on day 4 post-infection, was used as read-out. The percentage of inhibition was calculated by subtracting background (number of fluorescent pixels in the untreated infected control wells) and normalizing to the untreated-uninfected control wells (also background subtracted). The 50% effective concentration (EC_50_, the concentration of compound required for fifty percent recovery of cell-induced fluorescence) was determined using logarithmic interpolation. These experiments were performed in the presence of the Pgp-inhibitor CP-100356 (0.5 μM) in order to limit compound efflux. This graph was created using Graphpad Prism 9.2.0. The boxes extend from the 25^th^ to 75^th^ percentiles while the whiskers indicate the minimal and maximal values. The numbers above the X-axis indicate the number of measurements for each condition. While we determined the EC_50_ of remdesivir, molnupiravir and nirmatrelvir on Omicron we did not determine the EC_50_ on all VOC for all compounds tested. Due to time constrains we only used historical data from our database for the other VOCs and thus some of the values are depicted as “ND” (“Not Determined”). For the same reason, we included EIDD-1931 to allow comparison with molnupiravir as both compounds are intracellularly converted to the same antiviral molecule and thus have the same EC_50_. The individual EC_50_s values of this study are available at Mendeley Data (doi: 10.17632/bmjc74dyjs.1).

The fact that these antivirals retain their activity on the different SARS-CoV-2 VOCs is in accordance with the observation that the target proteins of these antivirals are highly conserved. For the RdRp there are only two amino acid changes (P323L in all VOCs and G671S in Delta; position 4715/5063 in ORF1ab or 314/662 in ORF1b respectively) when compared with the ancestral lineage (NC_045512). As these are distant from the active site, a different susceptibility towards remdesivir or molnupiravir is not to be expected. For the Mpro there are also two amino acid changes described (K90R in Beta and P132H in Omicron; position 3353 and 3395 in ORF1ab respectively). Alike for the RdRp, these mutations are not located near the active site of the Mpro and hence no difference in susceptibility for nirmatrelvir is expected.

These results indicate that when more VOCs arise, due to antigenic drift, there is a high probability that they will remain sensitive towards current (and likely also future) antivirals that do not target the spike. It is therefore of utmost importance to develop more pan-corona antivirals as they will be an essential armor and complement vaccines in the strategy to control the current pandemic [23].

## Acknowledgements

We thank Lize Cuypers, Guy Baele, Simon Dellicour and the Belgian Genomic Surveillance program which contributed to the early detection of Omicron in Belgium. We thank Tina Van Buyten, Birgit Voeten, Joost Schepers, Thibault Francken, Kim Donckers, and Niels Cremers for excellent technical assistance. We thank Fran Berlioz-Seux, Betsy Russel and Rob Jordan for helpful discussion. Part of this research work was performed using the ‘Caps-It’ research infrastructure (project ZW13-02) that was financially supported by the Hercules Foundation and Rega Foundation, KU Leuven. This project has received funding from the European Health Emergency preparedness and Response Authority (HERA), the Covid-19-Fund KU Leuven/UZ Leuven and the COVID-19 call of FWO (G0G4820N), the European Union’s Horizon 2020 research and innovation program under grant agreements No 101003627 (SCORE project) and Bill & Melinda Gates Foundation (BGMF) under grant agreement INV-00636 and the Innovative Medicines Initiative 2 Joint Undertaking (JU) under grant agreement No 101005077. The JU receives support from the European Union’s Horizon 2020 research and innovation program, EFPIA, the Bill & Melinda Gates Foundation, the Global Health Drug Discovery Institute, and the University of Dundee.

## Notes

https://data.mendeley.com/datasets/bmjc74dyjs/1

### Competing Interest Statement

The authors have declared no competing interest.

### Summary of Updates

improvements of the text

